# Sex Differences in Behavioral Responses during a Conditioned Flight Paradigm

**DOI:** 10.1101/2019.12.20.885038

**Authors:** Chandrashekhar D. Borkar, Mariia Dorofeikova, Quan-Son Eric Le, Rithvik Vutukuri, Catherine Vo, Daniel Hereford, Alexis Resendez, Samhita Basavanhalli, Natalia Sifnugel, Jonathan P. Fadok

**Affiliations:** Department of Psychology, Tulane University, New Orleans, Louisiana 70118, USA; Tulane Brain Institute, Tulane University, New Orleans, Louisiana 70118, USA; Program in Neuroscience, Tulane University, New Orleans, Louisiana 70118, USA

**Author notes:** Corresponding author*: Jonathan P. Fadok, Tulane Brain Institute, Tulane University #1345, 6823 St. Charles Ave., New Orleans, LA 70118-5698.

**Keywords:** sex differences, defensive behavior, fear conditioning

## Abstract

Females exhibit greater susceptibility to trauma- and stress-related disorders compared to males; therefore, it is imperative to study sex differences in the mode and magnitude of defensive responses in the face of threat. To test for sex differences in defensive behavior, we used a modified Pavlovian fear conditioning paradigm that elicits clear transitions between freezing and flight behaviors within individual subjects. Female mice subjected to this paradigm exhibited higher percentages of freezing behavior compared to males, especially during the intertrial interval period. Female mice also exhibited more cued freezing in response to the conditioned stimuli in the last block of extinction training. Furthermore, there were sex differences in the expression of other adaptive behaviors during fear conditioning. Assaying rearing, grooming, and tail rattling behaviors during the conditioned flight paradigm yielded measurable differences across sessions and between males and females. Overall, these results provide insight into sex-dependent alterations in mouse behavior induced by fear conditioning.

**Highlights:**

- Male and female mice do not differ in conditioned flight behavior.
- Female mice exhibit more freezing behavior.
- Rearing, self-grooming, and tail rattling behavior changes across days.
- Male mice exhibit more rearing and grooming behavior.
- Female mice exhibit more tail rattling behavior.

## 1. Introduction

Women have an approximately twofold higher risk of developing post-traumatic stress disorder (PTSD) compared to men [1,2], and they also display greater vulnerability to developing anxiety-related disorders [3]. Maladaptive behavioral responses to threats are associated with trauma- and anxiety-related disorders, making it imperative to study sex differences in the mode and magnitude of defensive behavior.

Pavlovian fear conditioning is a powerful model system that has provided tremendous insight into the neural mechanisms underlying fear-related learning and memory, mechanisms that are likely dysregulated in PTSD and anxiety disorders. Although a majority of research employing Pavlovian fear conditioning has used exclusively male subjects, some studies have explored sex differences in fear-related learning and memory. For example, female mice have been shown to generalize contextual fear more than males [4], while other studies have found that female mice and rats have impaired cued fear extinction [5–7]. Although these investigations provide insight into sex differences in generalization and extinction learning, they have often categorized freezing as the sole measurable index of fear.

According to the predatory imminence theory, different defensive behaviors are elicited depending on threat intensity, proximity, and context [8]. Based on this theory, we recently developed a modified Pavlovian conditioning paradigm that elicits clear transitions between conditioned freezing and flight behavior within individual subjects [9]. Therein, the conditioned flight paradigm was used to reveal important amygdala circuitry for the selection of defensive behavior; however, it utilized exclusively male mice. Therefore, we aimed to test the extent of sex differences in the conditioned flight paradigm. Additionally, we set out to examine sex differences in other adaptive behavioral changes in response to fear conditioning.

## 2. Materials and Methods

### 2.1 Animals

Adult male and female C57/BL6J mice (Jackson laboratory, USA) aged 3-5 months were used for the present study. All mice were individually housed on a 12 h light/dark cycle throughout the study with *ad libitum* access to food and water. Behavioral experiments were performed during the light cycle. All animal procedures were performed in accordance with institutional guidelines and were approved by the Institutional Animal Care & Use Committee of Tulane University.

### 2.2 Conditioned flight paradigm

Two different contexts were used for the conditioned flight paradigm. Context A consisted of a clear cylindrical chamber with a smooth floor, while Context B consisted of a clear square enclosure with an electrical grid floor (Med Associates, Inc.) used to deliver alternating current footshocks (ENV-414S, Med Associates Inc.). These two chambers were cleaned with 1% acetic acid and 70% ethanol, respectively. An overhead speaker (ENV-224AM, Med Associates, Inc.) was mounted above the chambers to deliver auditory stimuli at 75 dB. A programmable audio generator (ANL-926, Med Associates, Inc.) generated auditory stimuli. Behavioral protocols were generated using MedPC software (Med Associates, Inc.) to control auditory stimuli and shock via TTL pulses with high temporal precision.

The study sessions were conducted over a period of 4 days. Day 1 (pre-exposure) included a 3 min baseline period followed by 4 presentations of a serial compound stimulus (SCS) of 20 sec total duration in context A, with an 80 sec average pseudorandom intertrial interval (ITI) (range 60-100 sec). The SCS was a serial presentation of 10 sec pure tone (500 ms, 7.5 kHz pips at 1 Hz) and 10 sec white noise (500 ms pips at 1 Hz). The white noise was random and composed of frequencies from 1–20,000 Hz. The pre-exposure session lasted for 590 sec in total. On Day 2 and Day 3 (conditioning), mice were subjected to Context B and, after a 3 min baseline period, presented with five pairings of the SCS co-terminating with a 1 sec, 0.9 mA AC footshock, with a 120 sec average pseudorandom ITI (range 90-150 sec). Each conditioning session lasted for 820 sec in total. Day 4 consisted of the extinction session, also conducted in Context B. Following a 3 min baseline period, mice were presented with 16 trials of the SCS without footshock, with a 90 sec average pseudorandom ITI range (60-120 sec) spread over a total period of 1910 sec. This paradigm was also described in detail in [9].

### 2.3 Quantification of behavior

During the study, subjects were recorded and analyzed using Cineplex software (Plexon). Contour tracking ensured reliable data on relative position, while a central computer synced event markers to their real-time occurrences (Cineplex Studio, Plexon). Videos were scored for freezing behavior by an observer blind to the experiment with a frame-by-frame analysis of pixel changes (Cineplex Editor, Plexon). By determining a calibration coefficient using the chambers’ known size and the camera’s pixel dimensions, speed (cm/sec) was extracted using the animal’s center of gravity. Jumping escape behaviors were scored manually from video files. Flight score was calculated by dividing the average speed during each CS by the average speed during the 10 sec pre-CS (baseline) and then adding 1 point for each escape jump (speed_CS_/speed_BL_ + # of jumps). A flight score of 1 therefore indicates no change in flight behavior from the pre-CS period. In addition, videos were scored for rearing, grooming intervals, and tail rattling behaviors (Cineplex Editor, Plexon). To compare these adaptive behaviors effectively between fear conditioning and extinction sessions, we separated the extinction recording data into the first 820 sec (early extinction) and last 820 sec (late extinction) periods in order to better detail behavioral transitions. We excluded two female mice from the analysis on Days 3 and 4 because they jumped out of the conditioning context in response to the white noise stimulus on Day 3, precluding an accurate analysis of behavior.

### 2.4 Statistical Analysis

Data were statistically analyzed using Prism 8 (GraphPad Software, Inc.). All data were checked for normal distribution using the Shapiro-Wilk normality test (α=0.05), and the appropriate parametric/non-parametric test for significance was run. No statistical methods were used to predetermine sample size.

## 3. Results

### 3.1 Comparison of male and female mice in the conditioned flight paradigm

We used a four-day conditioned flight paradigm as previously described (Figure 1; [9]). On Day 1, mice freely explored a novel environment (Context A) and a serial compound stimulus (SCS; see Methods) was presented four times (Figure 1a, b). On Days 2 and 3, mice were placed into the conditioning context (Context B) and presented with five pairings of the SCS co-terminating with a 1 sec, 0.9 mA electrical foot shock (Figure 1a; see Methods). Finally, on Day 4, extinction training occurred in Context B.

**Figure 1.**
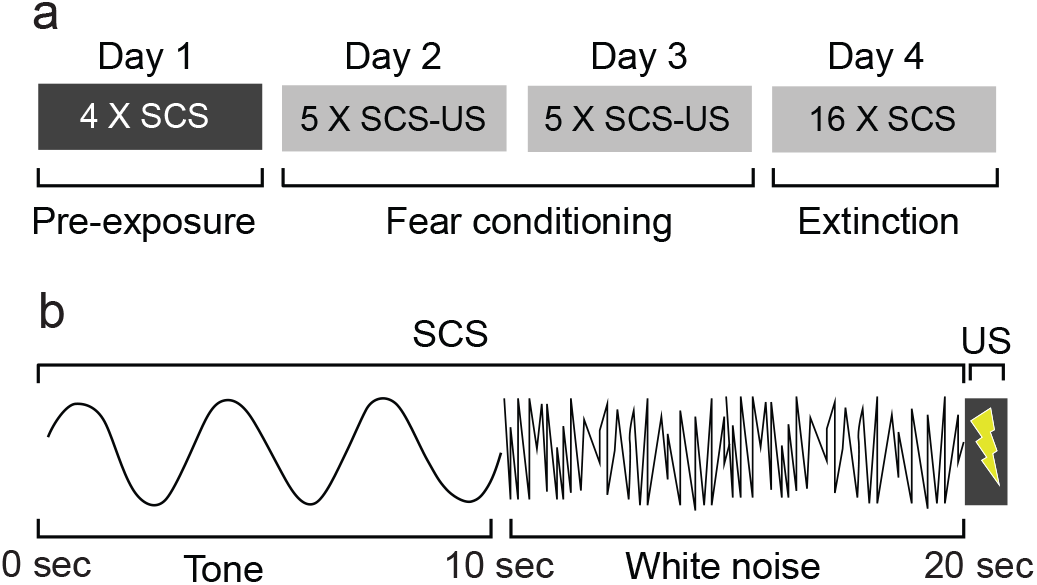
The conditioned flight paradigm. **a)** Diagram detailing the stages of the conditioned flight paradigm. Note that mice received footshocks (US) only on Days 2 and 3. **b)** Diagram detailing the composition of the SCS, as well as the timing of the US.

To analyze sex differences in conditioned freezing and flight behavior, we conditioned 10 male and 10 female mice in the flight paradigm. Both male and female mice developed robust freezing responses to the tone stimulus as well as robust flight responses to the white noise stimulus over the course of conditioning (Figure 2a-b). Consistent with our previous results [9], there was a significant cue X trial effect on flight behavior (Figure 2a; mixed-effects analysis, F_(1.849, 30.43)_ = 4.3, p<0.05). There was also a significant cue X trial effect on freezing behavior in both males and females (Figure 2b; mixed-effects analysis, F_(6.134, 101.0)_ = 18.48, p<0.0001).

**Figure 2.**
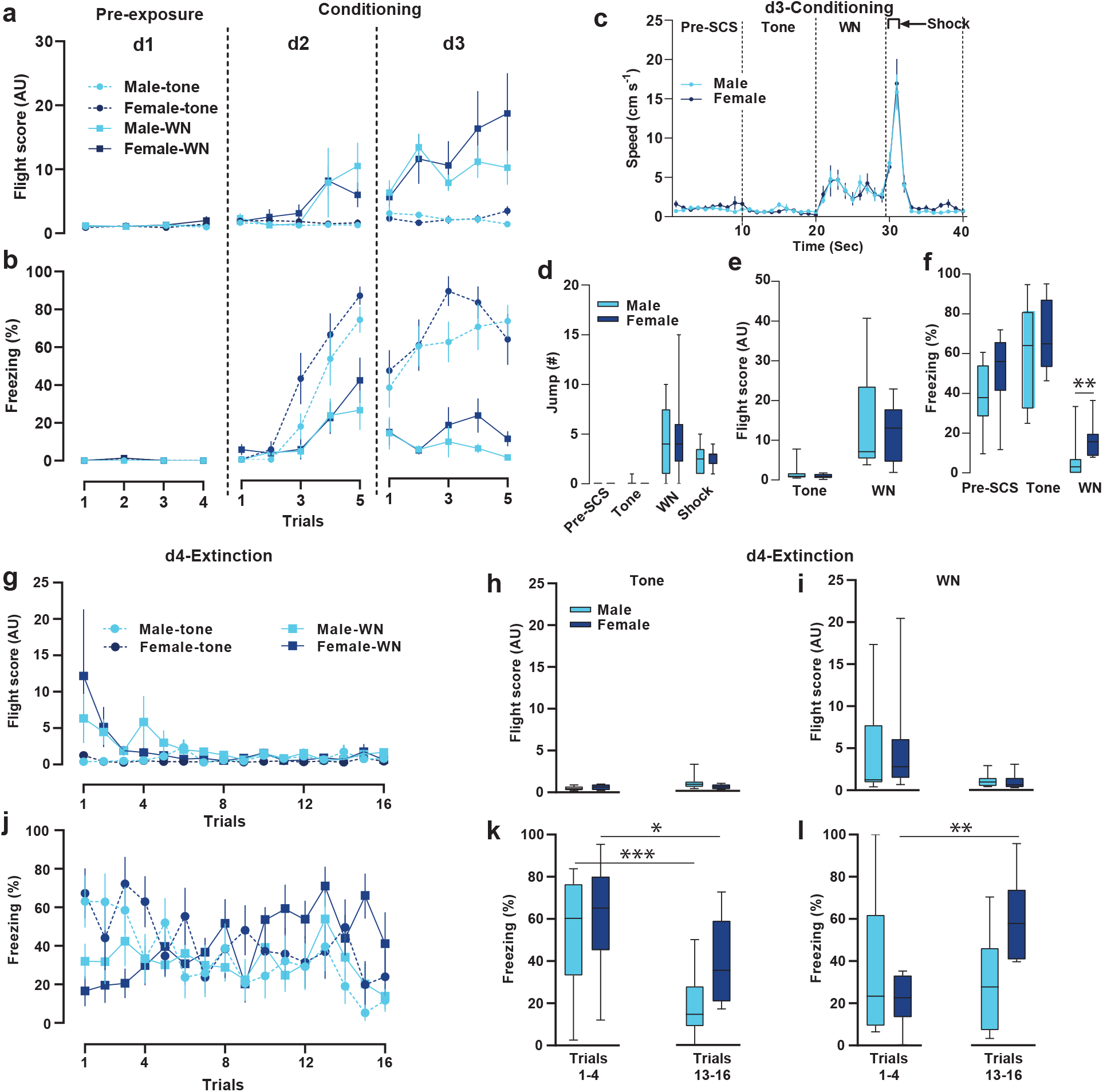
Sex differences in the conditioned flight paradigm. **a)** Average flight scores in males (n=10) and females (n=8) during the tone and white noise (WN) periods across Days 1-3. **b)** Average % freezing in males and females during the tone and WN periods across Days 1-3. **c)** Average speed (cm/s) of males and females during the pre-SCS, tone, WN, and shock periods on Day 3. **d)** Comparison of number of jump escape responses in males and females during the pre-SCS, tone, WN, and shock periods on Day 3. **e)** Comparison of flight scores in males and females in during the tone and WN on Day 3. **f)** Comparison of % freezing in males and females during the Pre-SCS, tone, and WN on Day 3. **g)** Average flight scores in males and females during the tone and WN on Day 4. **h)** Comparison of flight scores in males and females during the tone between early extinction and late extinction. **i)** Comparison of flight scores in males and females during the WN between early extinction and late extinction. **j)** Average % freezing in males and females during the tone and WN on Day 4. **k)** Comparison of % freezing in males and females during the tone between early extinction and late extinction. **l)** Comparison of % freezing in males and females during the WN between early extinction and late extinction. The represented values are means ± SEM. *p<0.05, **p<0.01, ***p <0.001.

Conditioned flight behavior is comprised of sudden increases in speed as well as escape jumps. There was no sex difference in white noise- or shock-induced speed increases on Day 3 of the flight paradigm (Figure 2c; two-way repeated-measures ANOVA, F_(39, 624)_ = 0.3171, p>0.99). There were also no differences in jumping behavior during the white noise (Figure 2d; average number of jumps: 5 for male and female; unpaired t test) nor in response to shock (average number of jumps: 3 for male and female, unpaired t test, p=0.97). Flight scores were significantly higher during the white noise for both males and females (Figure 2e; two-way repeated-measures ANOVA, effect of cue, F_(1, 16)_ = 21.18, p<0.001), and there was no significant sex difference in white noise-elicited flight (flight scores: 13.4 male and 12.1 female; Mann Whitney test, p=0.90).

Male and female mice exhibited contextual freezing on Day 3 (Figure 2f; pre-SCS: 39% versus 51%; unpaired t test, p=0.16) and elevated freezing during the tone period (60% versus 68%, unpaired t test, p=0.46). Males and females exhibited reduced freezing during the white noise stimulus (6.13% male, 16.25% female), consistent with the behavioral switch to flight. Freezing in female mice was significantly higher than in males during the white noise period (Mann-Whitney test, p<0.01).

On Day 4, mice underwent extinction training in the conditioning context. Flight behavior underwent rapid extinction in male and female mice (Figure 2g). During extinction training, flight responses were elicited specifically by the white noise stimulus and flight only occurred during the early extinction trials in both males and females (Figure 2h,i). There was no significant difference in flight scores between males and females (flight scores during first four trials: 4.63 male, 5.16 female; Mann Whitney test, p=0.23). Freezing behavior also changed over the course of extinction training (Figure 2j). There was significant within-session extinction in tone-induced freezing in male and female mice (Figure 2k; first four trials versus last four trials; Paired t test, p<0.001 male, p<0.05 female). Female mice had higher levels of freezing to the tone stimulus in the last bin of extinction trials compared to males, but this did not reach statistical significance (18.9% male, 32.6% female, unpaired t test, p=0.13). Freezing during the white noise period did not change during the extinction session for male mice (34.9% versus 30.3%; Paired t test, p=0.68); however, there was a significant increase in freezing to the white noise in female mice (Figure 2l; 1st four trials versus last four trial; 21.7% versus 55.6%; Paired t test, p<0.01).

### 3.2 Total freezing levels are higher in female mice

We next investigated sex differences in other dynamics of freezing and flight such as total duration, number of bouts, and duration per bout. On Day 3, there was no significant difference between males and females in the total duration of flight (Figure 3a; 12.2 sec male versus 14.2 sec female, unpaired t test, p=0.54), the number of flight bouts (Figure 3b; 7.4 male versus 7.8 female, unpaired t test, p=0.82), or the duration of flight bouts (Figure 3c; 1.7 sec male versus 1.8 sec female, unpaired t test, p=0.44). Interestingly, females had significantly higher total freezing duration on Day 3 (Figure 3d; 297.4 sec male versus 411.2 sec female, unpaired t test, p<0.05). Although there was no significant difference in the number of freezing bouts (Figure 3e; 67.4 male versus 61.8 female, unpaired t test, p=0.53), the average duration of individual freezing bouts was significantly higher in females (Figure 3f; 4.5 sec males versus 6.8 sec females, unpaired t test, p<0.05). The differences in total session freezing behavior between males and females can be attributed to significant differences in the levels of freezing during the ITI. Females expressed significantly elevated freezing during the ITI, both in total ITI freezing (Figure 3g; 264.6 sec male versus 372.7 sec female, unpaired t test, p<0.05) and in the duration of individual ITI freezing bouts (Figure 3h; 4.4 sec males versus 6.6 sec female, unpaired t test, p<0.05).

**Figure 3.**
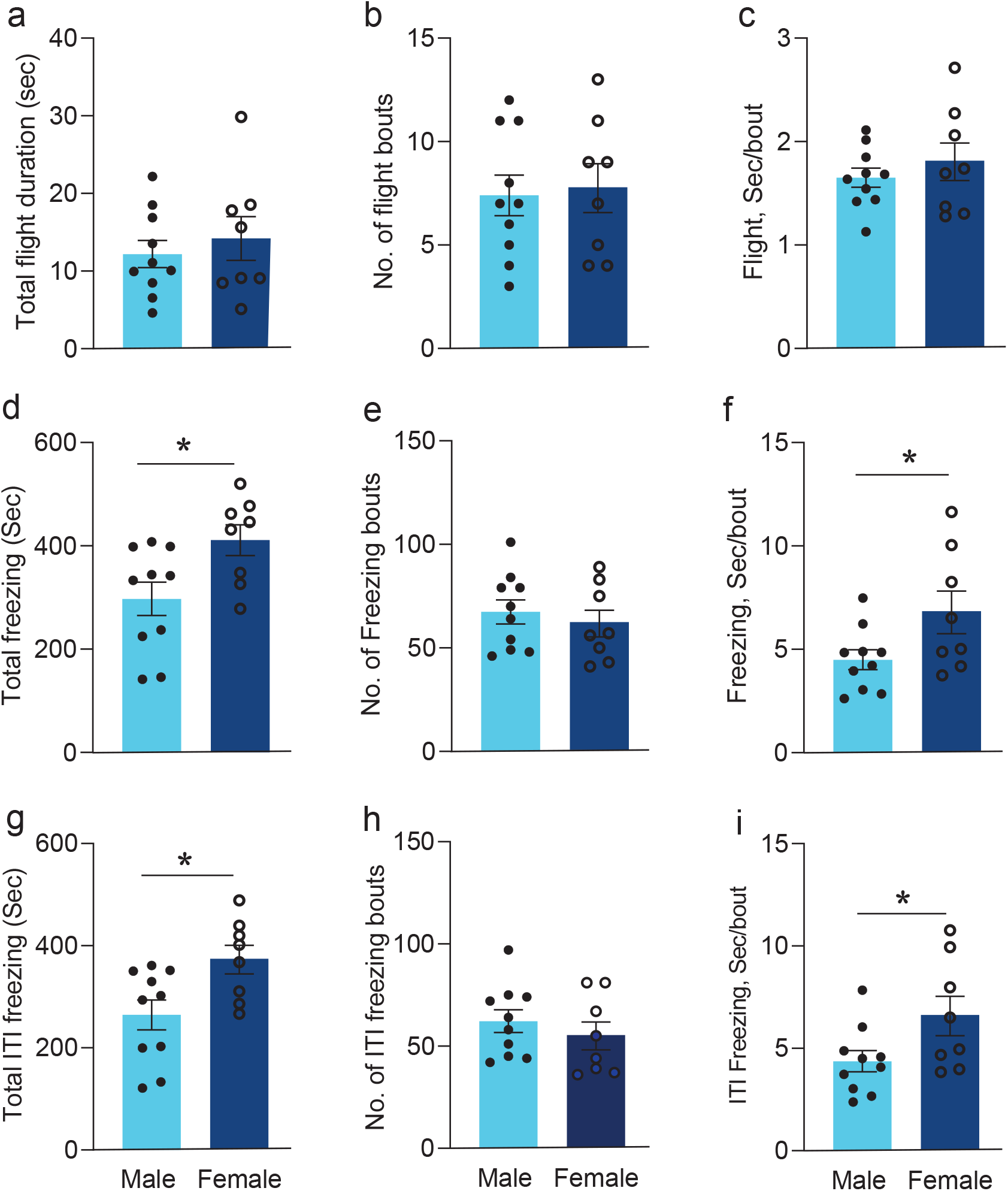
Comparison of freezing behavior between sexes on Day 3. **a-c)** Comparison of flight scores in males and females on Day 3 in terms of total flight duration **(a)**, number of bouts **(b)**, and duration of flight per bout **(c)**. **d-f)** Comparison of freezing in males and females on Day 3 in terms of total freezing **(d)**, number of bouts **(e)**, and duration of freezing per bout **(f). g-i)** Comparison of freezing in males and females during ITI on Day 3 in terms of total ITI freezing **(g)**, number of bouts **(h)**, and duration of freezing per bout **(i)**. Bars indicate means ± SEM. *p<0.05.

### 3.3 Changes in other adaptive behaviors during conditioning

In addition to flight and freezing, we quantified a number of other adaptive behaviors expressed during the conditioned flight paradigm. These included rearing, grooming, tail rattling, and general exploration. To demonstrate how these behaviors changed over trials, we generated mouse ethograms (Figure 4) to demonstrate behavioral dynamics over the course of conditioning.

**Figure 4.**
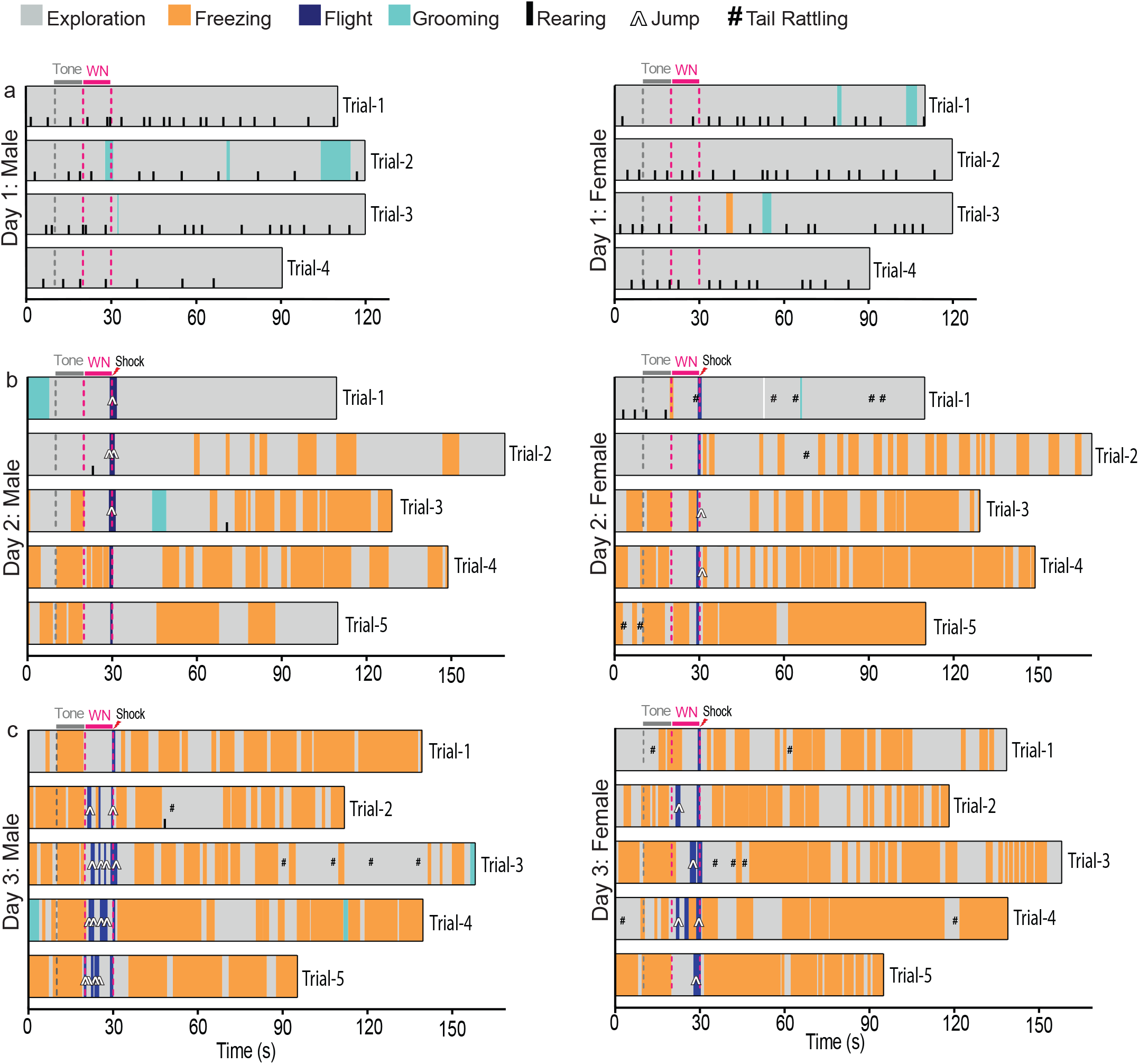
Representative ethograms of a male and female mouse during conditioning. Representative ethogram of males and females on Day 1 (pre-exposure) **(a)**, Day 2 (conditioning) **(b)** and Day 3 (conditioning) **(c)** illustrating the onset and duration of behaviors over trials in seconds. Legend differentiates recorded behaviors.

Rearing behavior changed significantly over the course of conditioning (Figure 5a; mixed-effects analysis, effect of session F_(2.119, 34.97)_ = 152.8, p<0.0001). Rearing behavior significantly decreased in male and female mice from Day 1 to Day 3 (Tukey’s multiple comparisons test; Day 1 versus Day 2, p<0.0001; Day 2 versus Day 3, p<0.0001). There was a significant increase in rearing during the extinction session compared to Day 3 (Tukey’s multiple comparisons test; Day 3 versus Day 4-early, p<0.001; Day 3 versus Day 4-late p<0.0001). Interestingly, males reared significantly more than females on Day 3 (Figure 5d; Mann-Whitney test, p<0.05).

**Figure 5.**
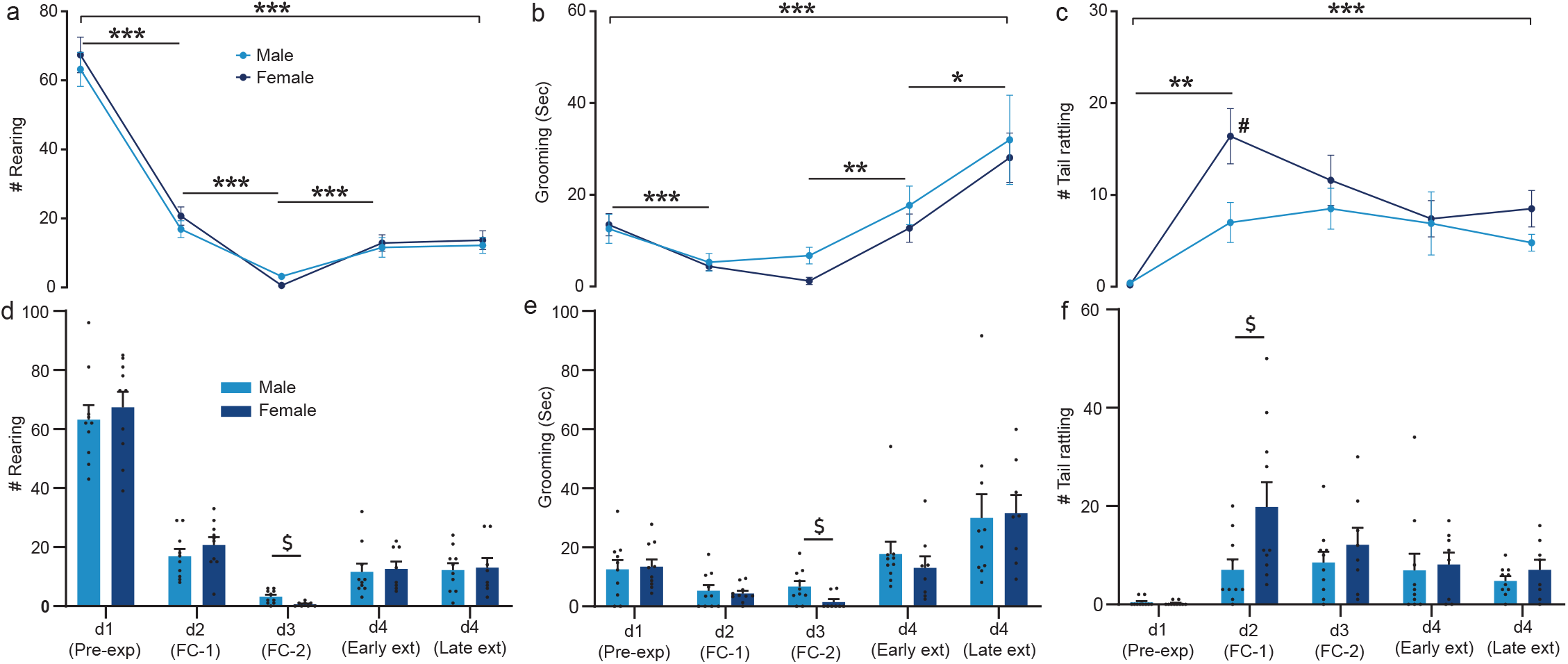
Changes in adaptive behaviors in the conditioned flight paradigm. Comparison of adaptive behaviors like rearing **(a,d)**, grooming **(b,e)**, and tail rattling **(c,f)** in males and females over all the experimental sessions of the conditioned flight paradigm. The represented values are means ± SEM. Mixed-effects analysis, effect of session ^*^p<0.05, ^**^p<0.01, ^***^p<0.001; ^#^p<0.05 male vs female. Mann-Whitney test, ^$^p<0.05.

Grooming behavior also changed significantly across sessions in males and females (Figure 5b; mixed-effects analysis, effect of session F_(1.853, 30.58)_ = 16.82, p<0.0001). Grooming decreased significantly from Day 1 to Day 2 (Tukey’s multiple comparisons test; p<0.001). Grooming increased over the course of the extinction session (Tukey’s multiple comparisons test; Day 3 versus Day 4-early, p<0.01; Day 4-early versus Day 4-late, p<0.05) eventually reaching levels that were significantly higher than those of the pre-conditioning session (Tukey’s multiple comparisons test; Day 1 versus Day 4-late, p<0.05). Just as with rearing, males groomed significantly more than females on Day 3 (Figure 5e; Mann-Whitney test, p<0.05).

Neither male nor female mice exhibited tail rattling behavior during the preconditioning session on Day 1 (Figure 5c). Occurrences of tail rattling increased for males and females during conditioning on Day 2 and remained elevated throughout Days 3 and 4 (Figure 5c; mixed-effects analysis, effect of session F_(2.090, 34.48)_ = 9.222, p<0.001; Tukey’s multiple comparisons test, Day 1 versus Day 2, p<0.01; Day 1 versus Day 3, p<0.001; Day 1 versus Day 4-early, p<0.05; Day 1 versus Day 4-late, p<0.001). Unlike the other behaviors, female mice tail-rattled significantly more than male mice on Day 2 (Figure 5f; unpaired t test, p<0.05).

## 4. Discussion

Using a fear conditioning paradigm that elicits both freezing and flight responses, we analyzed sex differences in defensive behavior (Figures 1–3). We found that male and female mice do not differ in conditioned flight behavior, yet freezing behavior is significantly greater in females (Figures 2, 3). Specifically, we found that females freeze more overall (Figure 3d,f) on Day 3, and that this increase is attributable to increased freezing during the white noise period (Figure 2f), with the greatest difference during the ITI (Figure 3g-i). There were also sex differences in other forms of adaptive behavior as mice underwent conditioning. By constructing ethograms, we were able to visualize the dynamics of these changes on a trial-by-trial basis (Figure 4). Rearing, grooming, and tail rattling behaviors all changed across sessions and there were sex differences in the expression of all three of these adaptive behaviors (Figure 5).

Although we did not find sex differences in flight scores, flight bout duration, or number of flight bouts, other studies have reported that a greater proportion of female rats engage in darting behavior [10,11]. These same studies found that female rats freeze less than males despite having a greater locomotor response to the shock [10]. Other studies also report lower contextual freezing in female rats [12–14], but similar levels of freezing to the auditory CS [13]. Interestingly, rats that engage in darting behavior express enhanced retention of extinction memory [11]. In this current study, we found that female mice freeze more than male mice on Day 3 in response to white noise and during the ITI period (Figures 2 and 3). Female mice also freeze significantly more in response to tone and white noise in the late extinction session (Figure 2). This is consistent with previous studies reporting impaired tone extinction memory in female mice and female rats [5–7]. Other studies in mice report that females are more likely to show generalized contextual fear [4]. Although results from different studies may seem disparate, it is important to appreciate that defensive behavior repertoires and situational response selection are likely to vary by species, strain, and sex. Careful behavioral analysis is therefore imperative for determining the specific strengths of each model for understanding the dysfunctions characteristic to PTSD and anxiety disorders [15–18].

Stress has profound effects on behavior, yet there is little knowledge of the sex differences in behavioral expression impacted by fear conditioning. Therefore, we examined the effects of conditioning and extinction training on rearing, grooming, and tail rattling in the mice subjected to the conditioned flight paradigm. Rearing is an index of arousal and general exploratory behavior in response to novelty [19,20], and this behavior changes during commonly used tests of anxiety-like behavior [21–23]. Our results show that both male and female mice decrease rearing behavior during the conditioned flight paradigm, and although rearing does not return to pre-conditioning levels, there is a significant return of rearing behavior during extinction training (Figure 5a). The severe reduction in rearing activity could be the result of the stress induced during conditioning, which might be reduced during fear extinction [21,23]. Interestingly, male mice rear more than females on Day 3 of the flight paradigm, which suggests that they might be more resilient to the deleterious effects of stress on behavior.

Grooming behavior is a self-maintenance behavior that is often used for stress reduction [24]. In response to stress, self-grooming behavior follows an inverted-U shaped curve, occurring spontaneously at low arousal levels, increasing during moderate arousal, and deteriorating during high-stress conditions that elicit freezing or flight responses [19,24,25]. Here, we find that grooming behavior significantly decreases during the conditioning phase of the flight paradigm in both male and female mice, while grooming eventually exceeds baseline levels by the end of the extinction session (Figure 5b). Interestingly, male mice groom more than females on Day 3, suggesting that the stress of undergoing fear conditioning differentially affects the subject according to sex. Reduced grooming in female mice may be suggestive of impaired stress coping strategies in females exposed to threatening situations.

Tail rattling is an aggressive defensive response elicited during social confrontation or in response to threat [26–28]. During the conditioned flight paradigm, tail rattling increases on the first day of conditioning and decreases during extinction for both male and female mice (Figure 5c). We also find that female mice exhibit significantly greater numbers of tail rattles than males on the first day of conditioning. Because tail rattling is elicited primarily in response to threat [28], this could have implications for understanding how trauma differentially affects expression of defensive aggression in males and females. For example, this may have translational relevance for understanding aspects of the hypervigilance response observed in patients with PTSD [17,18].

Our findings point to differential regulation of defensive adaptive behaviors between males and females. Subsequent studies can build upon these findings to reveal sex differences in the circuits controlling bodily responses to stress, anxiety, and fear. For example, our previous work in male mice demonstrated that central amygdala circuits are essential for switching between freezing and flight responses [9]. That study and others have shown that somatostatin-expressing neurons of the central amygdala are necessary for freezing behavior [9,29], whereas neurons expressing corticotropin-releasing factor are necessary for flight [9]. Because females express higher levels of freezing than males, future studies should investigate potential sex differences in the function of central amygdala neuronal networks, especially in light of recent work that has described sex differences in other nuclei important for defensive behavior [30,31].

## Acknowledgements

Conceptualization, C.D.B. and J.P.F.; Methodology, C.D.B. and J.P.F.; Formal Analysis, C.D.B., M.D., Q.E.L., R.V., C.V., D.H., A.R., S.B., N.S.; Investigation, C.D.B. and Q.E.L.; Resources, J.P.F.; Writing – Original Draft, C.D.B., M.D., and J.P.F.; Visualization, C.D.B., D.H., and J.P.F.; Supervision, J.P.F.; Project Administration, C.D.B. and J.P.F.; Funding Acquisition, J.P.F. We thank all authors for their thoughtful comments on the manuscript. This work was supported by a Louisiana Board of Regents Research Competitiveness Subprogram award (104A-18).

